# rarestR: An R package using rarefaction metrics to estimate α-diversity (species richness) and β-diversity (species shared) for incomplete samples

**DOI:** 10.1101/2024.04.29.591713

**Authors:** Yi Zou, Peng Zhao, Naicheng Wu, Jiangshan Lai, Pedro R. Peres-Neto, Jan C. Axmacher

## Abstract

Species abundance data is commonly used to study biodiversity patterns. In this context, estimating α- and β-diversity based on incomplete samples can lead to ‘undersampling biases’. It is therefore essential to employ methods that enable accurate comparisons of α- and β-diversity across varying sample sizes. This involves relying on biodiversity measures that are focused on accurately estimating the total number of species within a community, as well as the total number of species shared by two communities. Rarefaction offers such a method, where α-diversity is estimated for standardized sample sizes. Rarefaction methods can also be used as a basis for β-diversity calculations for standardized sample sizes. In this application note, we introduce a new R package, `rarestR`, designed to estimate abundance-based α- and β-diversity measures for inconsistent samples using rarefaction metrics. Additionally, the package offers parametric extrapolations to estimate the total expected number of species within a single community and the total expected number of species shared between two communities. Furthermore, it provides visualization for the curve fitting associated with these estimators. Overall, the `rarestR` package is useful in estimating α- and β-diversity values for incomplete samples, for example in studies involving highly mobile or species-rich taxa. These species estimators offer a complementary approach to non-parametric methods, such as the Chao series of estimators.

## Introduction

Measures of biodiversity can be split into two primary dimensions: α-diversity and β-diversity. Alpha-diversity usually describes the species richness encountered within a community, while β-diversity describes change in species composition between communities. Note, however, that other definitions (e.g. α-diversity describing the degree of entropy, and β-diversity describing the ratio between regional (γ) and local diversity) are also applied (Jost 2006, Tuomisto 2010, Whittaker 1960, Whittaker 1972).

Biodiversity studies commonly rely on abundance-based data, where each individual represents a sampling unit, or on incidence-based data where observational events such as the number of traps or quadrats represent the sampling units (Chao and Chiu 2016). A common issue encountered in biodiversity assessments is ‘undersampling bias’, which occurs due to low sample completeness, i.e., a lower number of species observed in a sample when compared to the available species pool of the sampled community (Coddington et al. 2009, Schroeder and Jenkins 2018). This bias impacts the estimations of both α- and β-diversity measures. To address this problem, it is critical to employ robust methods of standardization when comparing α- and β-diversity across incomplete samples (Beck et al. 2013, Coddington et al. 2009). Furthermore, biodiversity measures also depend on accurate estimates of the total number of species within a community, and of the total number of species shared between two communities derived from incomplete samples. Different types of data require different approaches to address ‘undersampling bias’ (Chao and Chiu 2016) and, in this context, we focus on abundance-based data.

Rarefaction calculates abundance-based probability distributions for standardized sample sizes, in turn providing the expected species (ES) richness for a given standardized sample size (Hurlbert 1971, Sanders 1968, Smith and Grassle 1977, Gotelli and Colwell 2001). It thus enables comparisons of α-diversity (as standardized species richness) among samples for a common standardized sample size. Building upon this concept, Grassle and Smith (1976) introduced a measure for the Expected number of Species Shared (ESS) between two samples. This index can be further standardized, allowing estimations of β-diversity (i.e., compositional dissimilarities between samples) for standardized sample sizes (Trueblood et al. 1994, Zou and Axmacher 2020, Zou 2021).

Employing asymptotic approximation to fit rarefaction curves for variable sample sizes, Zou et al. (2023) have introduced a parametric index to estimate the Total Expected Species richness (TES) within a community. Building upon the same mathematical principles, fitting the ESS curve also enables the estimation of the Total Expected number of Species Shared (TESS) by two communities (Zou and Axmacher 2021).

This application note introduces a new R package, `rarestR`, designed to calculate both, abundance-based α- and β-diversity measures for incomplete and inconsistent samples based on rarefaction metrics. Additionally, it integrates parametric extrapolation tools to estimate the total species within a single community and the total number of shared species between two communities, based on incomplete samples - the TES and TESS values mentioned above. The package furthermore supports the visualization of curve-fitting for the respective estimators.

## Mathematical description

### ES, ESS, NESS and CNESS

Hurlbert (1971) introduced the concept of an Expected Species richness (ES) for randomly drawn subsamples of *m* individuals from a larger sample, based on a hypergeometric distribution, referred to as *ESa* (eq. 1).

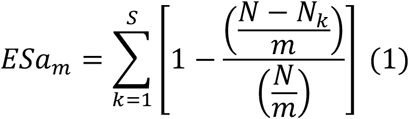

where *S* represents the number of observed species in the sample, *N* stands for the total sampled number of individuals, *N*_*k*_ denotes the number of individuals for species *k*, and *m* represents the standardized subsample size.

For communities containing an infinite number of individuals, Smith and Grassle (1977) proposed that the Expected Species richness follows a multinomial distribution, referred to as *ESb* (eq. 2). The results of *ESb* is linked to Simpson’s index at *m = 2* (i.e. ESb_2_ = Simpson + 1).

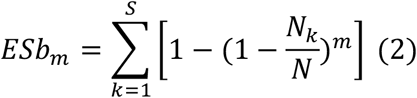

Building upon *ESa*, Grassle and Smith (1976) proposed a measure for the Expected Species Shared (ESS) between two communities. This concept once again involves drawing *m* individuals randomly from each sample, again assuming a hypergeometric distribution, with parameters varying according to sample properties (eq. 3):

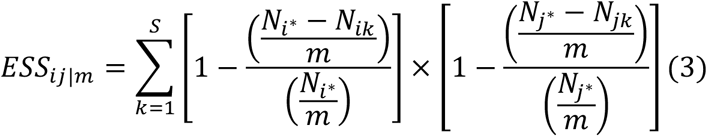

where *N*_*i\**_ and *N*_*j\**_ represent the number of individuals in the samples representing sites *i* and *j*, respectively; and *N*_*ik*_ and *N*_*jk*_ the number of individuals for species *k* in site *i* and *j*, respectively. Grassle and Smith (1976) also proposed an amended form of *ESS* based on *ESb*, but this has seldom been used.

Grassle and Smith (1976) additionally introduced a normalization of the *ESS* index based on the arithmetic mean, resulting in a distance measure with values ranging from 0 to 1, known as Normalized Expected Species Shared (NESS, eq. 4). The value of NESS is identical to the Horn-Morisita index (Morisita 1959) at *m = 1* (i.e. Horn-Morisita = 1 – NESS_ij|2_).

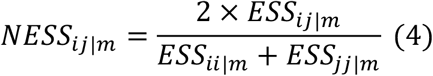

Trueblood et al. (1994) further modified this index, based on the geometric mean, termed the Chord-Normalized Expected Species Shared (CNESS, eq. 5).

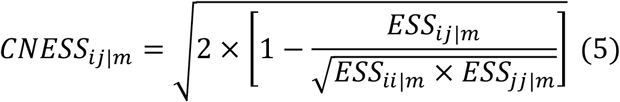

While CNESS has values ranging between 0 and 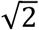, making it somewhat incompatible with other dissimilarity measures, Zou and Axmacher (2020) showed that the function can be modified by removing the 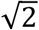 multiplier, resulting in a measure named CNESS_a_ (eq. 6).

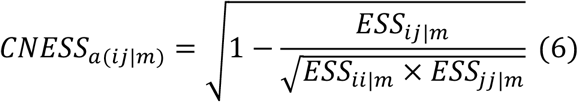

### TES and TESS

Zou et al. (2023) proposed curve-fitting for the relationship between the rarefaction curve (ES) and the standardized sample size (*m*), either based on a 4-parameter Weibull model (eq. 7) or a 3-parameter logistic model (eq. 8) (i.e., the Weibull-logistic model)

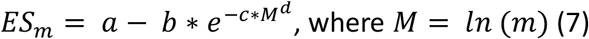

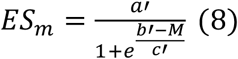

where a and a’ are the horizontal intercept values of the curve asymptotes that represent the Total Estimated Species richness (TES). The variance of this value is the estimated standard deviation (σ), as suggested by O’Hara (2005):

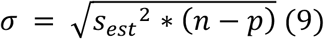

where *s*_*est*_ represents the standard error of TES obtained from the model fit, *p* the number of estimated parameters (i.e., either 3 or 4), and *n* the number of knots used in the model fitting. As ES has two different mathematical expressions, *ESa* and *ESb*, TES can be estimated separately from these two different models, resulting in TESa and TESb. The mean value of TESa and TESb, named ‘TESab’, provides a third measure, with a variance of:

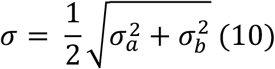

where *σ*_*a*_ and *σ*_*b*_ are the standard deviations of TESa and TESb based on formula 9.

Similarly, the relationship between ESS and the standardized sample size *m* can be fitted using the Weibull-logistic model, as proposed by Zou and Axmacher (2021). This allows the estimation of a Total number of Expected Species Shared (TESS) between two communities, with the variance estimated according to eq. 9.

## Package overview

The `rarestR` package can be downloaded and installed from CRAN (https://cran.r-project.org/web/packages/rarestR/index.html) or from GitHub (https://anonymous.4open.science/r/rarestr). The package contains four main functions, namely `es()`, `ess()`, `tes()`and `tess()`, and a training dataset, `share.` TES and TESS can be visualized via the `plot_tes()` and `plot_tess()` functions, or directly using the `plot()` function. Here are the descriptions of these functions:

1. `es(x, m, method, MARGIN)`calculates the rarefied number of species based on ‘expected species richness’ measures. The input `x` is a vector or a matrix representing the number of individuals for each species in one (vector) or across multiple sites (matrix). Parameter `m` represents the standardized subsample size (number of individuals randomly drawn from the sample), which by default is set to *m*=1, but can be varied according to users’ requirements. For *ESa*, `m` cannot be larger than the sample size. Argument `method` is the estimation approach of Expected Species used, with two options “a” and “b” available to calculate *ESa* and *ESb*, respectively, with the default set as “a”, returning identical values to the `rarefy()` function in the “`vegan`” package (Oksanen et al. 2018), but without providing a standard error. Argument `MARGIN` is a vector giving the subscripts which the function will be applied over, inherited from the `apply()`function.
2. `ess(x, m, index)`estimates the similarity between two samples based on the Expected Species Shared (ESS)-measure, using abundance data for the species contained in each sample. The input `x` is a community data matrix (sample x species; samples representing local communities), of which the sample name is the row name of the matrix. Argument `m` is the standardized sample size, again by default set to *m*=1. Rows with a total sample size <*m* will be excluded automatically from the analysis. Parameter `index` is the distance measure used in the calculation, as one of the four options “CNESSa” (formula 6), “CNESS” (formula 5), “NESS” (formula 4) and “ESS” (formula 3), with the default set as “CNESSa”. The function returns a pairwise distance matrix.
3. `tes(x, knots)`calculates Total Expected Species (TES) base on TESa, TESb, and their average value TESab. The input `x` is a data vector representing number of individuals for each species. Argument `knots` specifies the number of separate sample sizes of increasing value used for the calculation of TES between 1 and the endpoint, which by default is set to `knots=40`. The function returns a list with a self-defined class “`rarestr`”, which contains a summary dataframe of the estimated values and their standard deviations based on TESa, TESb, and TESab, and the detailed results of the models used in the estimation of TES, either ‘logistic’ or ‘Weibull’.
4. `tess(x, knots)`calculates the number of Total Expected Species Shared (TESS) between two samples. The input `x` is a data matrix for two samples representing two communities. Argument `knots` has the same meaning and default value as that in the function `tes().` The function returns a list with the self-defined class “`rarestr`”, which contains a summary dataframe of the estimated values and their standard deviations of TESS, and the detailed results of the model used in the estimation of TES, either ‘logistic’ or ‘Weibull’.
5. `plot_tes(x)` and `plot_tess(x)`visualize the fitted curve of the models for calculating TES and TESS, respectively. The input `x` is an object with the “`rarestr`” class (i.e. an object returned by the `tes()` or `tess()`function). Alternatively, and more user-friendly, the fitted curves can be plotted via the generic `plot()` function, which automatically evokes an appropriate visualization function.

The package contains a dataset named `“share”`, comprising three samples randomly drawn from three simulated communities. Every community consists of 100 species with approximately 100,000 individuals, following a log-normal distribution (mean = 6.5, SD = 1). Setting the first community as reference (i.e., fully randomly generated), the second and third community share a total of 25 and 50 randomly selected species with the reference community. A detailed description of the reference and scenario communities can be found in Zou and Axmacher (2021). The `share` dataset represents a random subsample of 100, 150 and 200 individuals, randomly drawn from these three communities, containing 58, 57 and 74 species, respectively.

## Performance of rarefaction-based α-diversity and β-diversity

Here we briefly tested the performance of rarefaction-based alpha and beta diversity that is available in the `rarestR` package using simulated data. We would like to emphasize that the application of different metrics depends on different scenarios. Comprehensive evaluations of the biodiversity metrics is beyond the scope of this application note (but see Beck and Schwanghart 2010, Zou and Axmacher 2020). The brief demonstration of its performance in contrast to different metrics aims to help users to recognize that these metrics can be accurate. It also offers examples for using the package effectively.

For alpha diversity, we tested the performance of ESa (Hurlbert rarefaction, eq. 1) and ESb (Smith and Grassle rarefaction, eq. 2) for its precision and accuracy in estimating species richness for samples with incomplete and inconsistent sizes, and how the performance changes with increased sample sizes. Two samples were randomly drawn from the simulated reference community (i.e. 100 species of 100,000 individuals, following a log-normal distribution). The first sample contained *n* individuals, while the second sample contained twice this original sample size (i.e., *2*n* individuals), with *n* increasing from 10 to 150 randomly drawn individuals. The ratio between two samples was calculated for ESa and ESb. We contrasted the performance of ESa and ESb with the Shannon diversity index, the (bias-corrected) Chao1 lower boundary species richness estimator (O’Hara 2005), and the observed species richness. We repeated the process 1000 times to obtain the mean and 95% quantile of the respective values.

Results show that, in comparison, ESa is accurate and precise in reflecting the true difference for samples with different sample sizes, even for very low sample sizes of *m*=10 individuals. In contrast, ESb, Shannon diversity and the Chao1 estimator all underestimate this difference, with Chao 1 having the lowest precision (Figure 1a). Observed species richness has the lowest accuracy, always underestimating the true difference.

**Figure 1.**
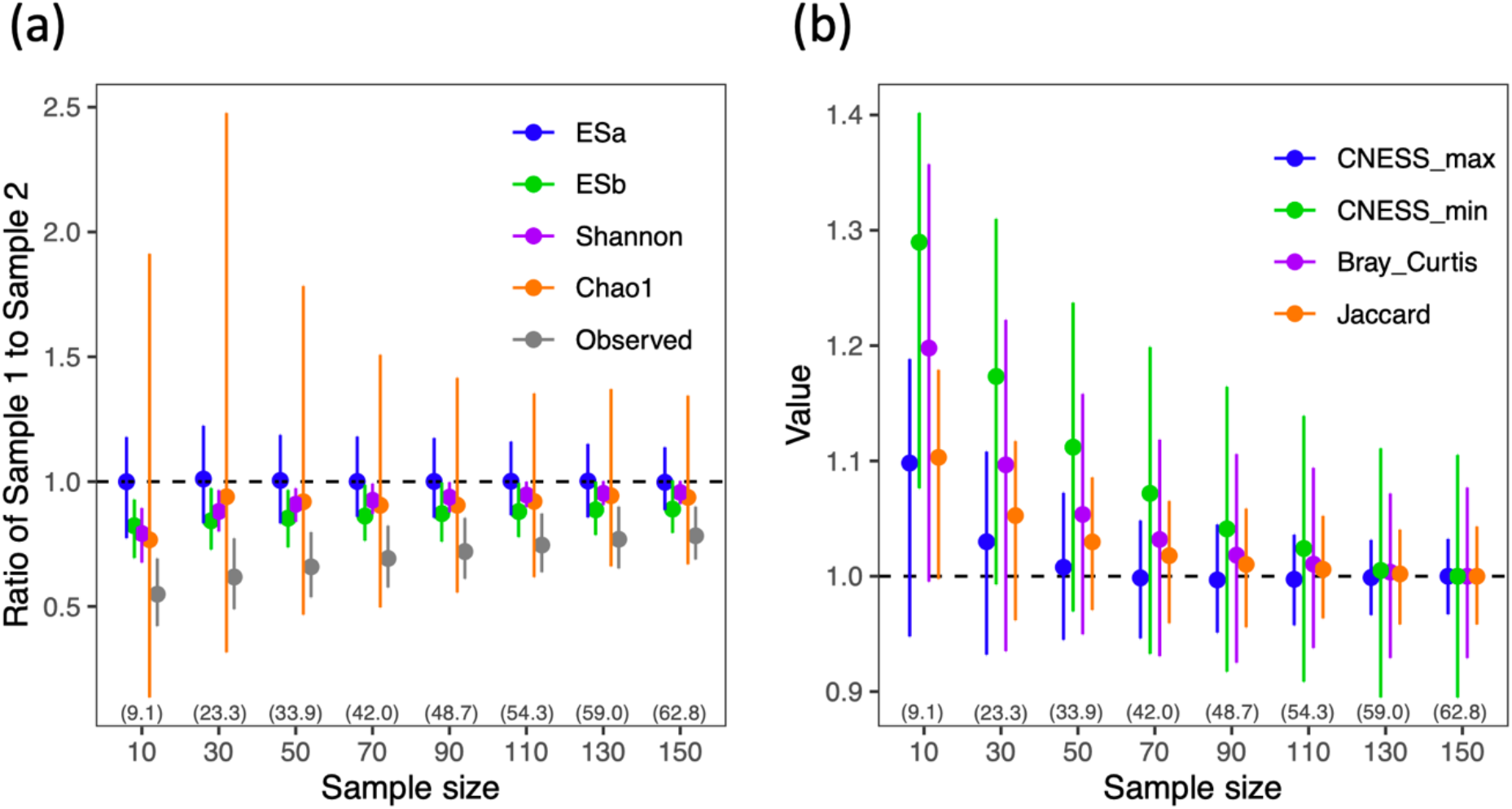
(a) The change of the ratio between sample 1 and sample 2 that are randomly collected from a communities (contain 100 species of approximately 100,000 individuals) for ESa, ESb, Shannon diversity, Chao1 value and observed species richness along with the increase of the sample size of sample 1 (sample 2 is twice large as sample 1); dashed line represents the actual ratio; values in brackets refer to the mean % sampling completeness as the proportion of the number of species sampled to the overall number of species in the pool reached for a specific sample size. (b) Ratio of the results from sample size *n* to its maximum (i.e. n = 150) for CNESS at the smallest m (CNESS_min, m = 1), largest m (CNESS_max, m = n), the Bray-Curtis index and the Jaccard index; two samples are randomly drawn from two communities (each contains 100 species of approximately 100,000 individuals) which shares a total of 25 species. Sample one drew n individuals from the first community and sample two drew 2*n individuals from the second community. Dots and error bars refer to mean and 95 quantile from 1000 repetitions for both cases.

For beta diversity, we tested the performance of CNESSa (e.q. 6) and NESS (e.q. 4) for two different sample sizes *m*— the minimum value (*m*=1) and the maximum value that can be selected (*m*= maximum common sample size across samples). Two samples were randomly drawn from the above-mentioned two communities, the first reference community and the second community that shared a total of 25 species with the reference. We randomly drew *n* individuals from the first community and *2*n* individuals from the second one, with *n* increasing from 10 to 150.

As the value of the ESS series depends on the parameter *m*, with a small *m* value emphasizing similarities in the composition of abundant species, while a large *m* value leading to estimations of similarities in the overall community (Zou and Axmacher 2020), accuracy cannot be accessed. Therefore, we focused in this instance on the ‘stability’ of the results from CNESS and NESS by calculating the ratio of the results for different sample sizes n. We compared our results with the commonly used Bray-Curtis (abundance-based) and Jaccard dissimilarity (incidence-based) indices. Again, we repeated the process 1000 times to obtain the mean and 95% quantile values.

Results show that CNESS at large *m* values has a higher stability and precision than both, Bray-Curtis and Jaccard indices, particularly when the sample size is larger than 50. CNESS at *m*=1 in contrast is relatively unstable and imprecise (Figure 1b). In comparison, NESS values have a very low precision for both, small and large values of *m*, while values for large *m* are again more stable than values for a small *m* (Appendix 1).

## Working examples

We demonstrate here the use of functions in the `rarestR` package by using simulated and empirical data. The simulated data was taken from the `“share”` file as described earlier. For the empirical data, we used the `mite` data provided by the `vegan` package (Oksanen et al. 2018). This data comprises 70 sites (communities) of 35 species of Oribatid mites. To improve the clarity of the results, we chose to analyze only the first 20 sites in the dataset (3447 individuals of 33 species).

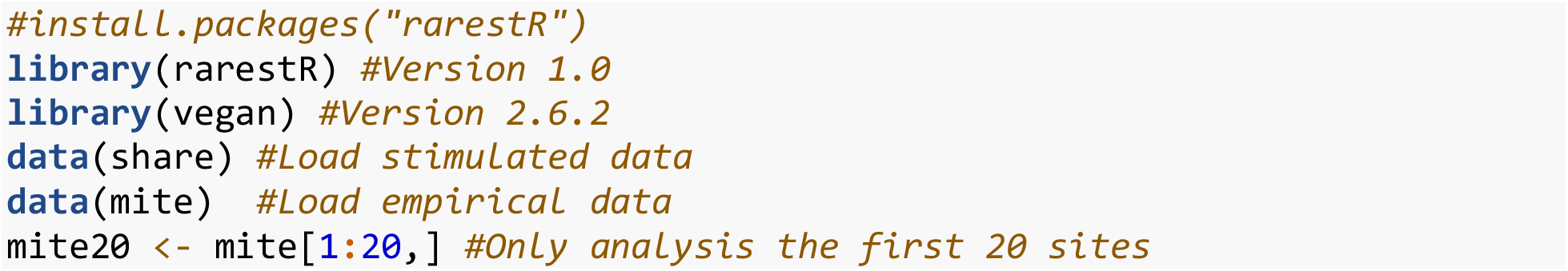

### Function es()

We demonstrate the application of ES for a maximum standardized value where no site (sample) is disregarded (*m* = 90, i.e., the minimum sample size across all sites). When *m* exceeds the total sample size for a given sample, “NA” will be returned by the software.

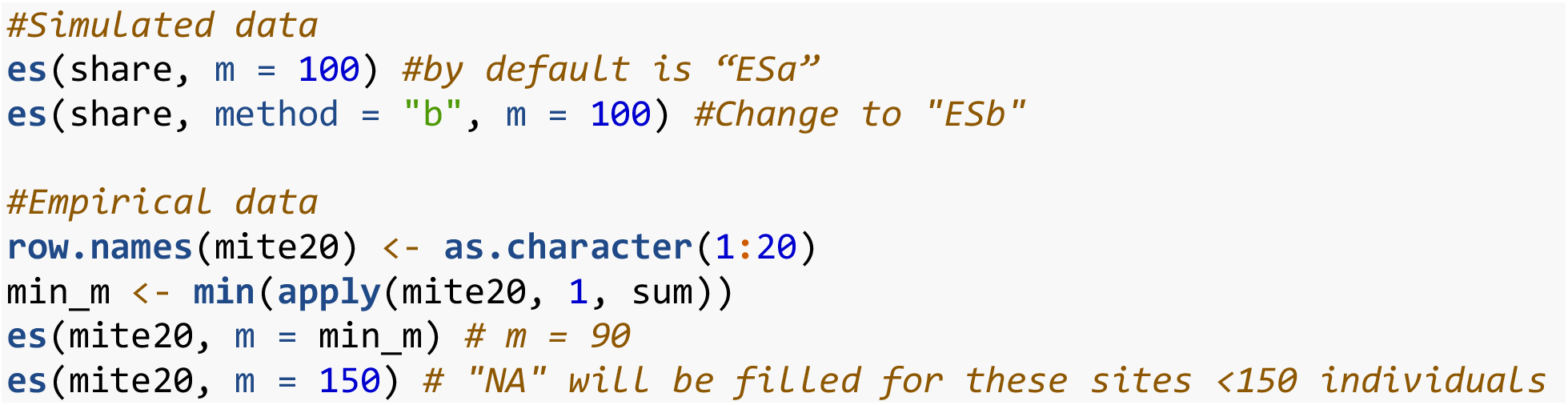

### Function ess()

For ESS measures, we calculated the minimum standardized value, *m*=1, and the maximum standardized value (*m* = 90) for CNESS and NESS measures. We then visualized the CNESS and NESS matrix results using non-metric multi-dimensional scaling (NMDS).

NMDS plots show diverging results for the two different standardized sample size values (*m*=1 and *m*=90) based on the CNESS dissimilarity matrices for the `mite` data. For this example, results indicate that species dissimilarity patterns are different when we focus on dominant species, or on the overall community composition. For instance, for dominant species (*m*=1), site 15 clusters together with several other sites, but for the overall community (*m*=90), it is positioned distinctly apart from all other sites (Figure 2). In contrast, NMDS plots based on the NESS matrices show rather homogeneous patterns for both small and large *m* values (Appendix 2).

**Figure 2.**
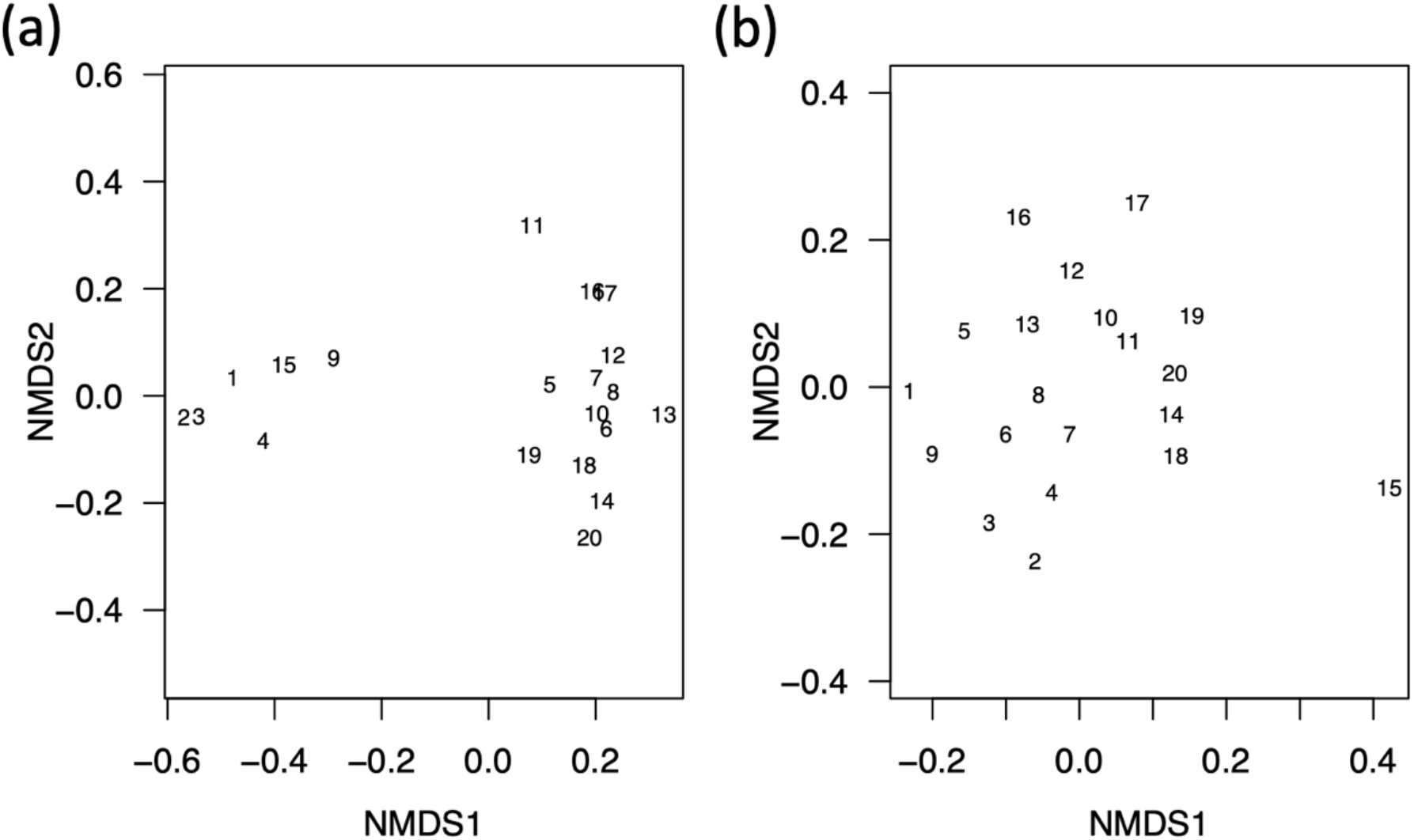
NMDS (non-metric multi-dimensional scaling) based on the CNESS (Chord-Normalized Expected Species Shared between two samples) dissimilarity measures for m =1 (a) and m = 90 (b) for `mite` data in the `vegan` package; numbers represent the site ID.

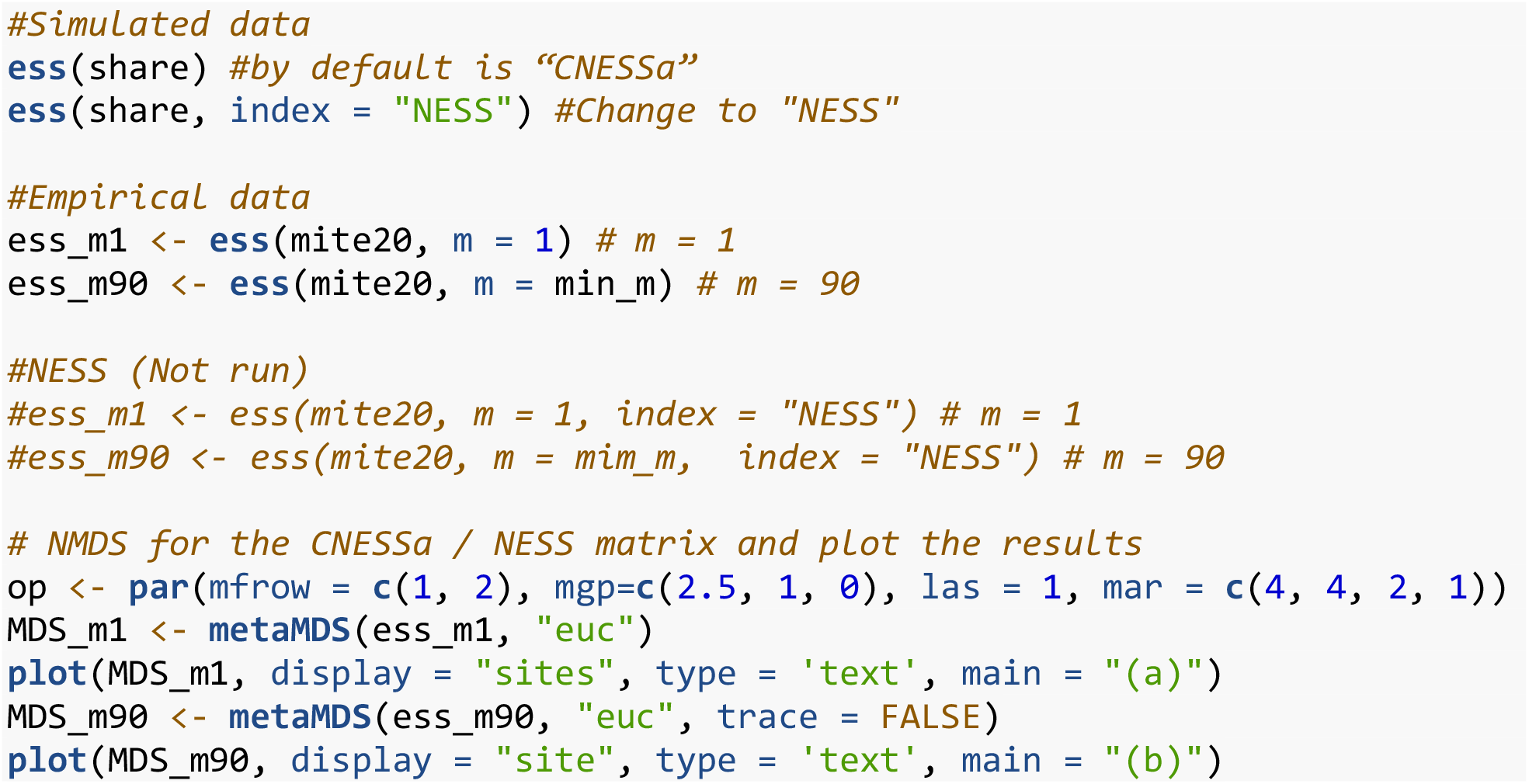

### Function tes()

TES results (i.e., based on curve-fitting) for the simulated data show TESa=138.5 and TESb=92.63 (Figure 3), with a TESab=115.56 for the first sample. For TES measures of the empirical data, we calculated the value for pooled data of the 20 sites, which contains 33 observed species. Results show TESa=24.63, TESb=34.14 and TESab=34.39, which is very close to the overall species riches in the `mite` data.

**Figure 3.**
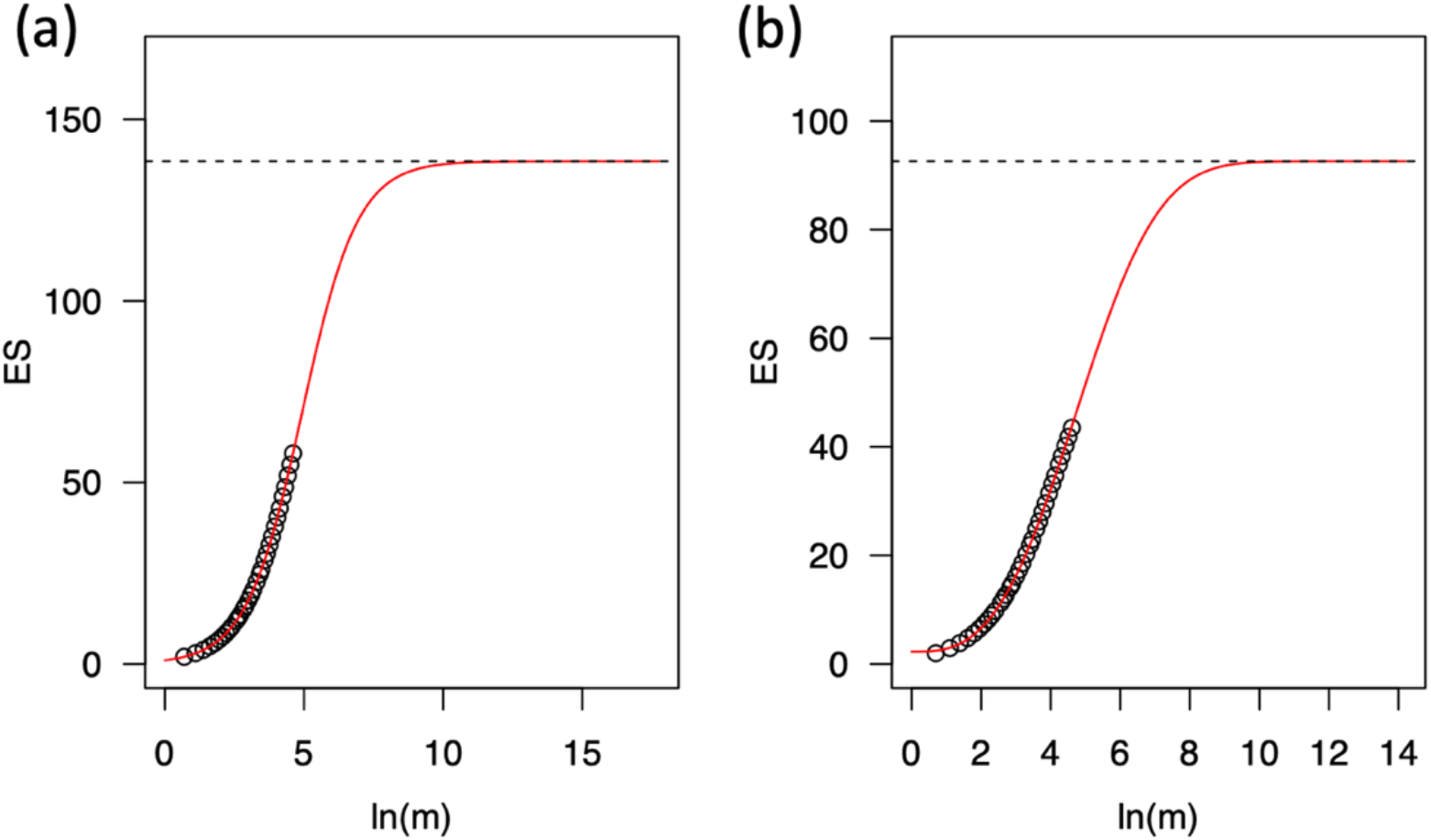
Number of expected species (ES) based on ESa (a) and ESb (b) estimations versus the standardized sample size (*m*) for the simulated data of ∼100,000 individuals split across 100 species; solid lines refer to the model fit; dashed lines refer to the total expected species (TES), i.e., the asymptotic value of the model.

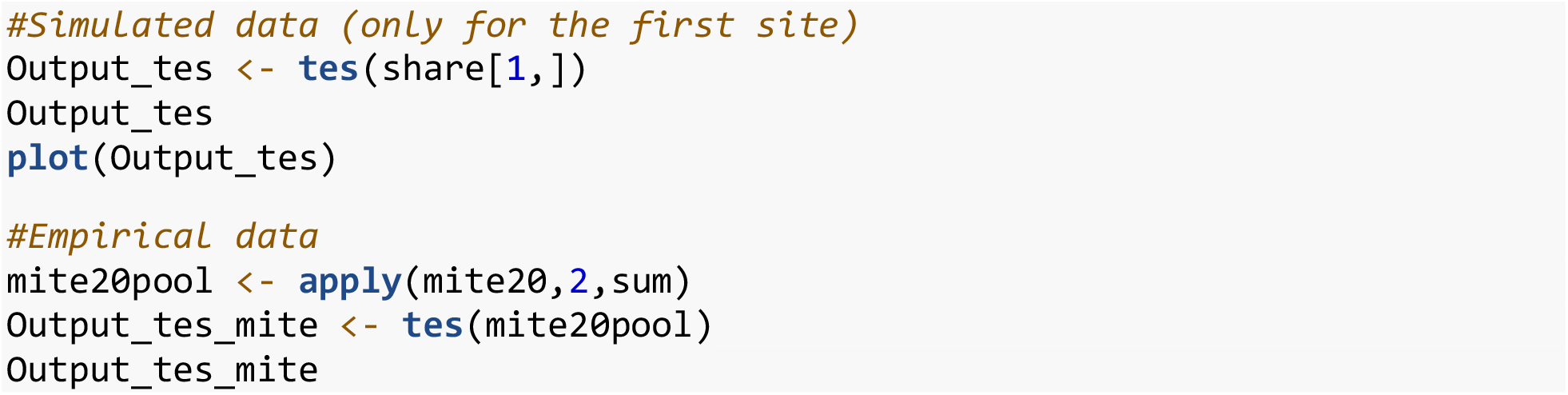

### Function tess()

The TESS value for the simulated data between the first and the second samples is 23.28, and between the first and third sample is 40.16 (Appendix 3a and b), which are relatively close approximations of the “real” values of 25 and 50 species. For the empirical data, in order to obtain a robust estimation, we calculated the estimated shared species with the pooled data of site 1 to site 10, and pooled data of site 11 to site 20. Results show that they are expected to share 32.14 species (Appendix 3c).

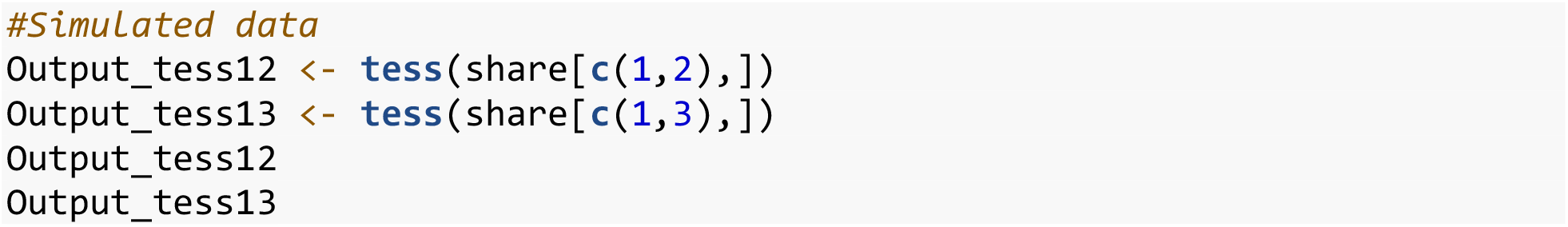
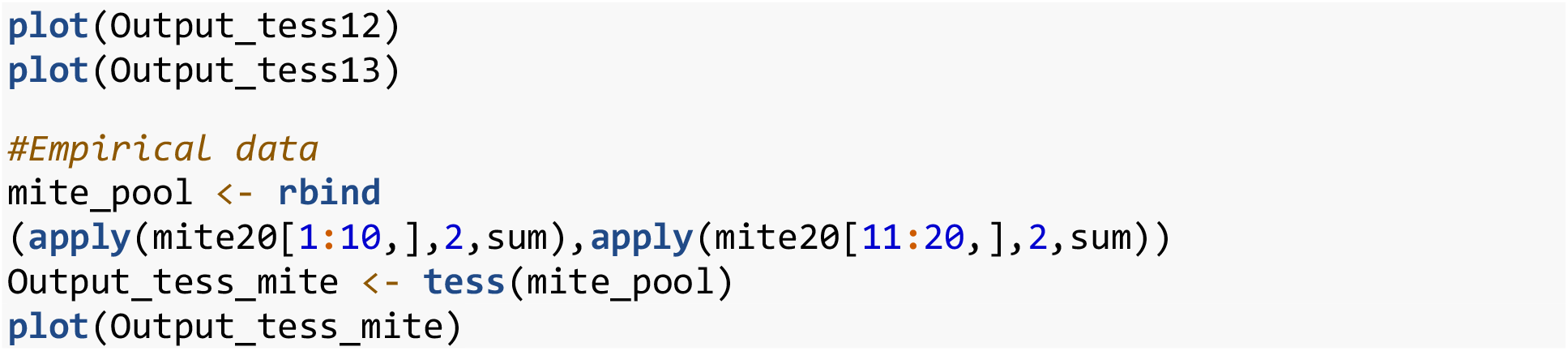

## Discussion

The `rarestR` package integrates calculations of the expected species number (i.e., rarefaction and extrapolation) for a single sample, and the expected number of species shared between communities represented by two samples, based on species abundance data. Both `tes()` and `tess()` functions use asymptotic approximation to extrapolate rarefaction curves, thereby estimating the total number of species within single communities and the total number of species shared by pairs of communities.

As mentioned above, the `es()`function calculates a rarefaction value, where the calculation of “ESa” is based on a hypergeometric distribution, which is identical to the `rarefy()` function from the `vegan` package (Oksanen et al. 2018). The calculation of “ESb” is based on a multinomial distribution that is not available as a choice in the `vegan` package. Simulation results suggest that “ESa” outperforms “ESb” in precision and accuracy in detecting the true difference of species richness for incomplete and inconsistent samples. It is important to note, however, that, where *m* exceeds the total sample size, the `es()`function returns “NA”. This behaviour differs slightly from the `rarefy()` function, which returns the number of observed species. This distinction is intentional, highlighting the importance of excluding samples larger than the standardized sample size from comparisons. Moreover, this approach aligns with the approach used in the β-diversity comparison performed by the `ess()`function.

The function `ess()` calculates the β-diversity based on an adjustable standardized sample size. Generally, β-diversity measures fall into two classes: direct calculation of the ratio between regional (γ) and local (α) diversity, and multivariate measures based on pairwise dissimilarities (Anderson et al. 2011). Our ESS-based β-diversity estimate fundamentally differs from the recently developed beta diversity rarefaction and extrapolation methods in the package `iNEXT.beta3D` (Chao et al. 2023), as well as from the sample coverage-based rarefaction β-diversity proposed by Engel et al. (2021). Both approaches estimate β-diversity based on the ratio between estimated γ- to α-diversity, providing an average (regional) measure of β-diversity across all communities. In contrast, our approach used in the `ess()` function (index “CNESSa”, “CNESS” and “NESS”) estimates β-diversity based on pairwise dissimilarities in communities represented by (incomplete) samples. In this context, the CNESS index for large *m* values is less sensitive to sample size variations compared to, for example, Bray-Curtis and Jaccard indices. The results from CNESS furthermore can vary when adjusting the value for the parameter *m* — a small value reflecting composition dissimilarities among dominant species, while larger values increasingly accounts for overall community similarities (Zou and Axmacher 2020). In contrast, the NESS index appears less precise and generates homogeneous results across varying *m* values when compared with CNESS. Therefore, we recommend the use of the CNESS index, and to select both small and large *m* values in order to comprehensively interpret the results and underlying community structures.

Our `tes()`extrapolation method differs from that in the `iNEXT` package, which estimates species richness (Hsieh et al. 2016), and its extension, `iNEXT.3D` (to phylogenetic and functional diversity, Chao et al. 2021). `iNEXT` generally employs an abundance-based method to estimate the number of species without asymptotic values (but note, asymptotic values may be achieved based on diversity measures, see Hsieh et al. 2016), which may yield infinite estimates for extensive sample sizes (Cayuela et al. 2015). The `tes()`function calculates the total expected species based on asymptotic parametric curve fittings. Unlike traditional curve-fitting approaches that depend on specific species abundance distributions (Walther and Moore 2005), TES provides flexibility and robust applicability across various species abundance distribution models (Zou et al. 2023). TES is hence comparable with non-parametric estimators such as Chao 1 and ACE (e.g., in `vegan` package) (Chao 1984, Chao and Lee 1992). Additionally, we provide visualizations for these estimations, offering a complementary approach to these non-parametric methods.

Our function `tess()`finally calculates the estimated number of shared species between two communities, again based on asymptotic parametric curve fitting. To our knowledge, the only other shared species richness estimators available are Chao1-shared and ACE-shared in the `SpadeR` package (Chao et al. 2016). However, TESS generally outperforms these in terms of precision and accuracy, especially when dealing with unequal sample sizes (Zou and Axmacher 2021). Similar to `tes()`, visualizations are available through the `plot()` function, allowing researchers to graphically interpret their curve fittings. Integrating TESS with other species richness estimators would enable users to more accurately estimate true species (dis)similarities, based on both shared and unique species numbers, as listed in Koleff et al. (2003). However, caution should be exercised when combining estimators, as this inadvertently leads to a reduced precision (Zou and Axmacher 2021).

Generally, species estimators, despite being commonly employed to accommodate varying sample sizes in the comparison of biodiversity across samples, have low precision particularly for relatively small sample sizes. Consequently, we do not recommend users to generally use such estimators for comparing multiple samples since their primary utility (in our view) lies in estimating sampling completeness within a given target community. Only where estimated completeness is high should the estimators be used to then also estimate true species richness and similarity values. This rationale underpins our decision to design the `tes()` function for a single sample, and the `tess()`function for two samples, only. For multiple sample comparisons accounting for different sample sizes, we recommend using the ES for α-diversity and CNESS for β-diversity. Therefore, both the `es()` and `ess()` functions can be applied for multiple samples (communities) in our package.

In summary, the `rarestR` package proves valuable for ecologists working on α- and β-diversity, especially when dealing with incomplete and inconsistent sample sizes, which is a prevalent characteristic in samples of ecological communities, but particularly in samples of highly mobile and species-rich taxa. Additionally, it provides visual estimations for species richness and the number of shared species between two communities based on individual samples, offering a complementary approach to non-parametric methods, such as the Chao series of estimators (Chao 1984, Chao et al. 2000, Chao et al. 2023).

## Supporting information

**Appendix 1.**
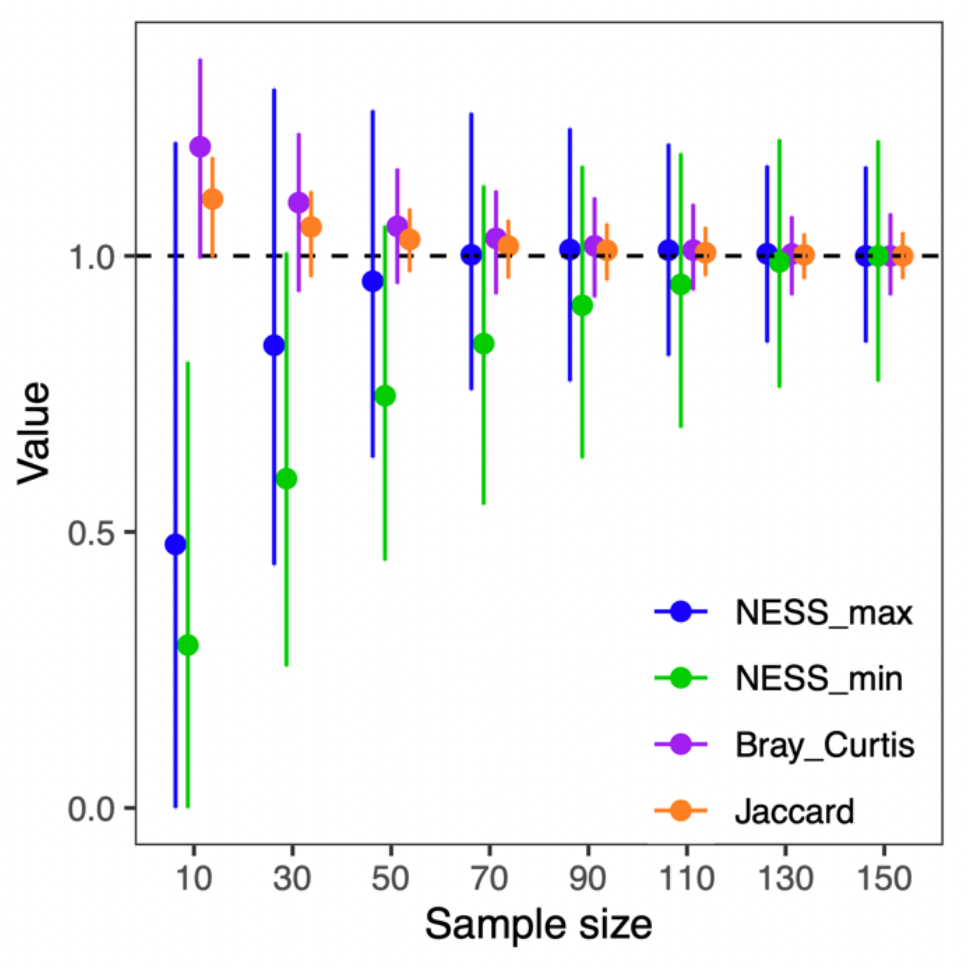
Ratio of the results from sample size *n* to its maximum (i.e. n = 150) for NESS at the smallest m (NESS_min, m = 1), largest m (NESS_max, m = n), the Bray-Curtis index and the Jaccard index; two samples are randomly drawn from two communities (each contains 100 species of approximately 100,000 individuals) which shares a total of 25 species. Sample one draws n individuals from the first community and sample two draws 2*n individuals from the second community. Dots and error bars refer to mean and 95 quantile from 1000 repetitions for both cases.

**Appendix 2.**
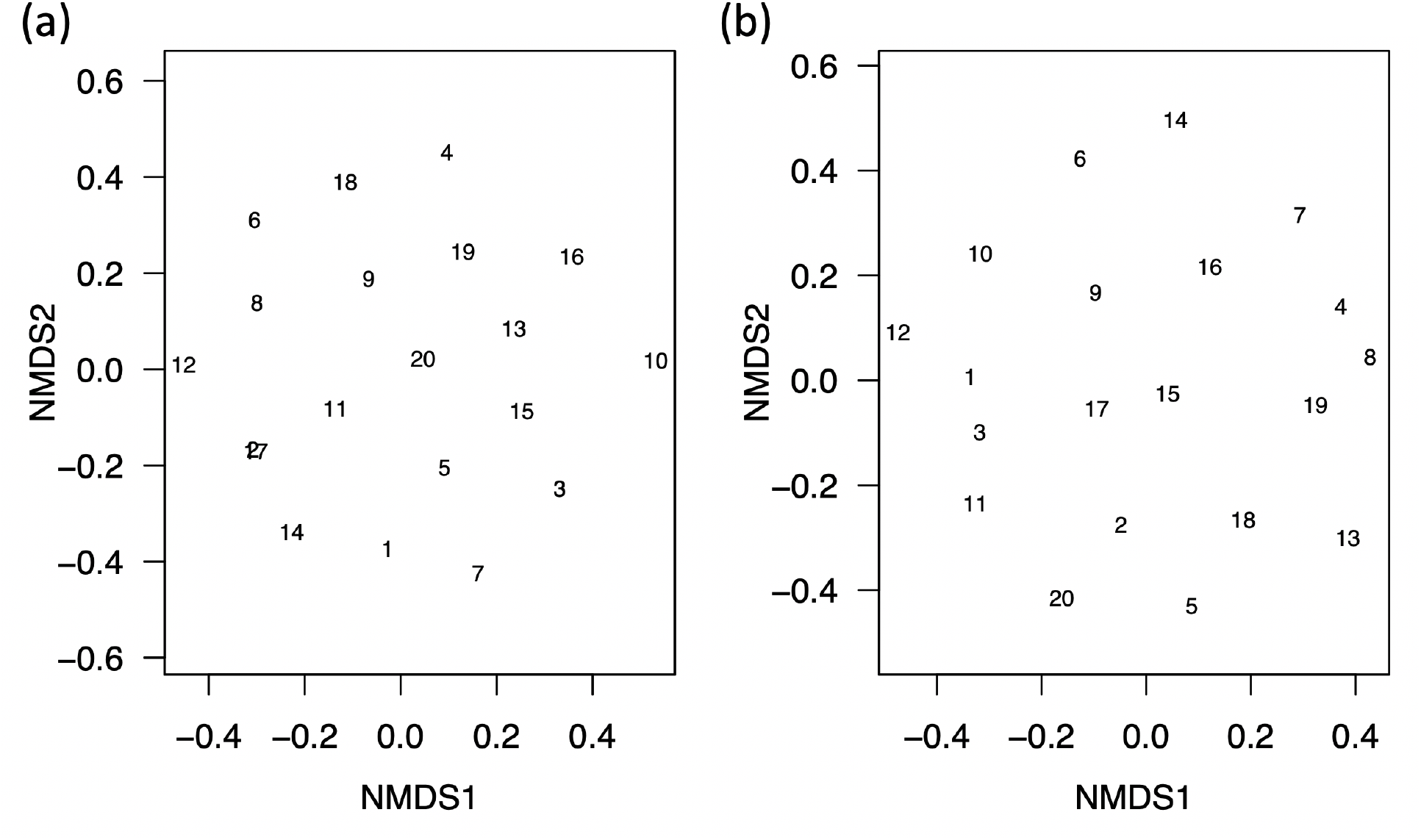
NMDS (nonmetric multi-dimensional scaling) based on the NESS (Normalized Expected Species Shared between two samples) dissimilarity measures for m =1 (a) and m = 90 (b) for `mite` data in the `vegan` package. Numbers represent the site ID.

**Appendix 3.**
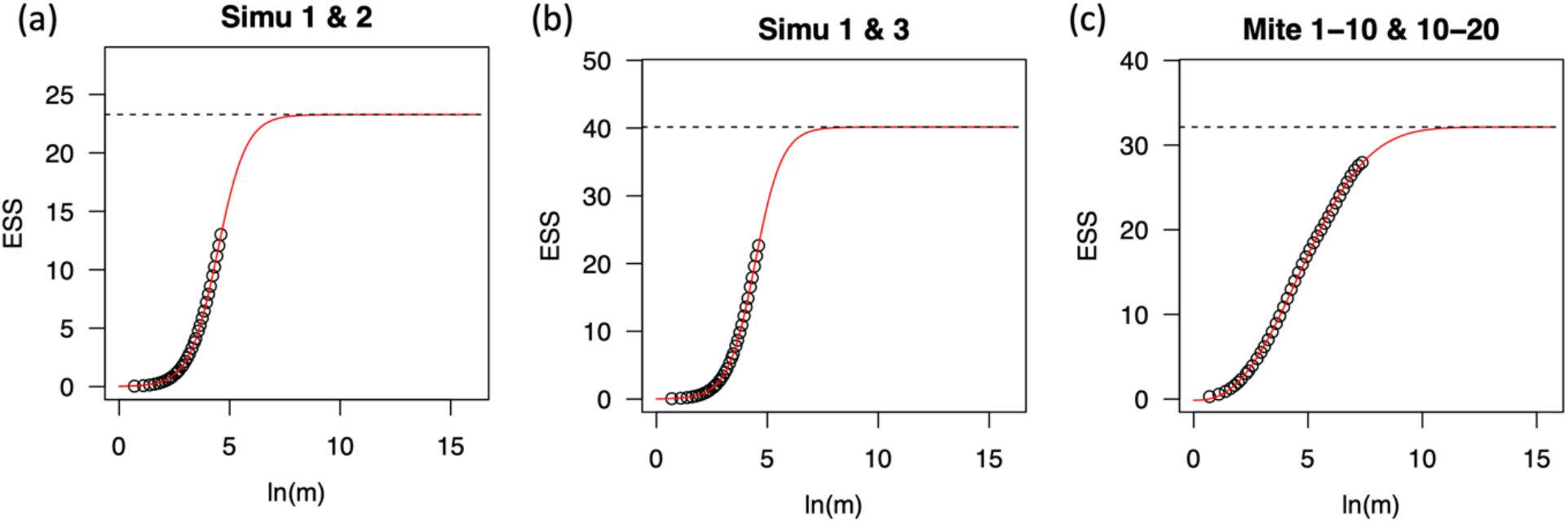
Number of expected species shared (EES) between the second (a) and third (b) sample with the control versus the standardized sample size (m) in the simulated `share` dataset, and for empirical data between pooling site 1 to 10 and pooling site 11 to 20 (c). Solid lines refer to the model fit; dashed lines refer to the total expected species shared (TESS) between two communities (i.e., asymptotic value of the model fit).

## Notes

**Conflict of interest** The authors declare no conflict of interest.

### Competing Interest Statement

The authors have declared no competing interest.

